# Prenatal benzene exposure alters offspring hypothalamic development predisposing to metabolic disease in later life

**DOI:** 10.1101/2023.01.05.522910

**Authors:** Lisa Koshko, Sydney Scofield, Lucas Debarba, Lukas Stilgenbauer, Mikaela Sacla, Patrick Fakhoury, Hashan Jayarathne, J. Eduardo Perez-Mojica, Ellen Griggs, Adelheid Lempradl, Marianna Sadagurski

## Abstract

The hypothalamus is essential in the regulation of metabolism, notably during critical windows of development. An abnormal hormonal and inflammatory milieu during development can trigger persistent changes in the function of hypothalamic circuits, leading to long-lasting effects on the body’s energy homeostasis and metabolism. We recently demonstrated that gestational exposure to benzene at smoking levels induces severe metabolic dysregulation in the offspring. Given the central role of the hypothalamus in metabolic control, we hypothesized that prenatal exposure to benzene impacts hypothalamic development, contributing to the adverse metabolic effects in the offspring. C57BL/6JB dams were exposed to benzene in the inhalation chambers exclusively during pregnancy (from E0.5 to E19). The transcriptome analysis of the offspring hypothalamus at postnatal day 21 (P21) revealed changes in genes related to metabolic regulation, inflammation, and neurodevelopment exclusively in benzene-exposed male offspring. Moreover, the hypothalamus of prenatally benzene-exposed male offspring displayed alterations in orexigenic and anorexigenic projections, impairments in leptin signaling, and increased microgliosis. Additional exposure to benzene during lactation did not promote further microgliosis or astrogliosis in the offspring, while the high-fat diet (HFD) challenge in adulthood exacerbated glucose metabolism and hypothalamic inflammation in benzene-exposed offspring of both sexes. These findings reveal the persistent impact of prenatal benzene exposure on hypothalamic circuits and neuroinflammation, predisposing the offspring to long-lasting metabolic health conditions.

## Introduction

Compelling evidence indicates that gestational exposure to various environmental toxins predisposes to metabolic disorders and Type 2 Diabetes (T2DM) later in life (1–3). Recent epidemiological studies have highlighted the association of VOC exposure and particularly benzene, one of the main volatile organic compounds (VOCs) in ambient air, with increased risk for gestational diabetes mellitus (GDM) (4). Indeed, increased offspring risk for cardiovascular and metabolic disease later in life has also been described in the context of the altered maternal environment, such as GDM, obesity, and exposure to environmental toxic compounds during pregnancy (5–7). In support, we have recently shown that gestational exposure to benzene at the levels relevant to cigarette smoking, induces deleterious effects on energy homeostasis, hyperglycemia, and insulin resistance in adult offspring (8). However, the mechanisms underlying fetal metabolic programming and predisposition to metabolic imbalance following gestational exposure to benzene remain unknown.

Benzene can cross the placenta, directly impacting gestation and triggering an inflammatory response in the fetus (9). The central nervous system (CNS) is particularly sensitive to environmental insults during development, and exposure to ambient concentrations of VOCs can have detrimental impacts on fetal brain development (10). One mechanism by which gestational exposure to VOCs may cause impaired health outcomes is via neuroinflammation mediated by the activation of glial cells (such as microglia and astrocytes), leading to adverse neural adaptations and neurotoxicity (11–14). Microglia, the major cell type responsible for CNS immune response, are exquisitely sensitive to stressors and are activated by inhaled components of urban air pollution through both direct and indirect pathways (15–17). Microglia regulate the innate immune functions of astrocytes through the release of interleukin 1 *α* (IL-1 *α*), TNF *α*, and C1q, inducing astrocytes to acquire a pro-inflammatory, and neuron-killing phenotype (18). Benzene exposure can induce microglia and astrocytes activation either directly, or via secreted inflammatory cytokines from the periphery that can cross the blood-brain barrier (BBB) (19).

The hypothalamus is one of the key regions of the CNS responsible for the regulation of energy homeostasis, satiety signals, and glucose metabolism (20, 21). As the rodent hypothalamus begins development early in gestation and continues postnatally (22, 23), both *in-utero* and postnatal environments can influence this process (22). We and others have shown that maternal under or overnutrition is critical for the offspring’s hypothalamic development, inducing hypothalamic inflammation and pro-inflammatory cytokines in the brain at fetal and postnatal stages (24–27). These developmental windows represent important intervals of vulnerability during which alterations in the maternal environment may lead to abnormal hypothalamic development and subsequent metabolic alterations.

In the present study, we aimed to elucidate the impact of prenatal benzene exposure (at smoking levels (8)) on the offspring’s hypothalamic development and in response to the diet-induced obesity (DIO) challenge. Our data provide concrete evidence for the sex-specific effect of prenatal benzene exposure on the hypothalamic transcriptome, metabolic pathways, and neuroinflammation that can subsequently act as a driver for late-life metabolic disease.

## Materials and Methods

### Animals

8-week-old C57BL/6 mice were obtained from The Jackson Laboratory. Mating of male and female mice began following 1-week habituation. Female mice were considered pregnant upon observation of a vaginal plug. Animals were kept on a 12-hour light cycle and had access to food and water *ad libitum*. C57BL/6 pregnant dams were exposed to benzene (50 ppm) for 6 hours per day (6h/day) from E0.5 to E19 using our FlexStream automated Perm Tube System (KIN-TEK Analytical, Inc) as previously described (19);(8). Offspring were fed HFD (45% kcal) at 22 weeks of age for 8 weeks. For leptin signaling, a separate cohort of 21-day-old offspring fasted for 4h followed by an intraperitoneal (i.p.) injection of leptin (5mg/kg) and perfused after 2h as done previously (28).

### RNA Extraction for RNA-sequencing

Before RNA extraction, tissue samples from the whole hypothalamus of P21 offspring were stored in RNA-later (Invitrogen, AM7020) at −80°C. The tissues were handled on dry ice, weighed, and immediately frozen with liquid nitrogen in screw cap tubes containing 3 mm Zirconium Beads (OPS Diagnostics, BAWZ 3000-300-23). 1 mL TRIzol Reagent (Life Technologies, 15596026) was added per 50-100 mg of tissue. Tubes were homogenized using the FastPrep-24 system on high for 30 seconds and visually observed for tissue disruption, performing additional homogenizations as needed. 200 ul chloroform (Sigma-Aldrich, 319988) was added, and samples were centrifuged at 12,000 x g for 15 minutes at 4°C. The upper aqueous phase was transferred to new Eppendorf tubes followed by RNA precipitation using 500 ul ice-cold isopropanol (Sigma-Aldrich, 190764) and 2 ul GlycoBlue (ThermoFisher, AM9516) for pellet visualization. Samples were then centrifuged at 12,000 x g for 15 minutes at 4°C. Supernatant was removed and discarded. The samples were washed twice with 75% ethanol (Fisher Scientific, 111000200), using a 5-minute 7,500 x g centrifugation each round for pellet collection. The pellet was air-dried for 10 minutes and resuspended in 20 ul RNase-free water (Life Technologies, AM9938). Samples were incubated at 60°C for 5 minutes, then stored at −80°C.

### Construction and Sequencing of Directional total RNA-seq Libraries

Libraries were prepared by the Van Andel Genomics Core from 500 ng of total RNA using the KAPA RNA HyperPrep Kit (Kapa Biosystems, Wilmington, MA USA). Ribosomal RNA material was reduced using the QIAseq FastSelect –rRNA HMR Kit (Qiagen, Germantown, MD, USA). RNA was sheared to 300-400 bp. Before PCR amplification, cDNA fragments were ligated to IDT for Illumina TruSeq UD Indexed adapters (Illumina Inc, San Diego CA, USA)). Quality and quantity of the finished libraries were assessed using a combination of Agilent DNA High Sensitivity chip (Agilent Technologies, Inc.), QuantiFluor® dsDNA System (Promega Corp., Madison, WI, USA), and Kapa Illumina Library Quantification qPCR assays (Kapa Biosystems). Individually indexed libraries were pooled and 50 bp, paired-end sequencing was performed on an Illumina NovaSeq6000 sequencer to an average depth of 30M raw paired reads per transcriptome. Base calling was done by Illumina RTA3 and the output of NCS was demultiplexed and converted to FastQ format with Illumina Bcl2fastq v1.9.0.

### RNA-sequencing data analysis

Adapter and low-quality sequences were trimmed from the reads using Trim Galore v0.6.0 with the ‘--paired’ parameter. Trimmed reads were aligned to the mm10 genome with GENCODE M24 gene annotations, using STAR v2.7.8a with the parameters ‘--twopassMode Basic’, and ‘--quantMode GeneCounts’ to generate the read counts per gene. Downstream analyses were conducted using the reverse-stranded counts produced by STAR. Transcripts with a total raw read count < 20 were filtered out from the analysis. The biological sex of the animals was confirmed using the expression of Xist. This led to the reassignment of samples OBFH14 and OBFH5 to the male and OBMH20 and OBMH4 to the female group, which is reflected in the metadata file. Data diagnostics identified and excluded outlier sample OBFH14 due to a > 2-fold higher Cook’s distance median compared with all other samples. Read count normalization and controls versus treatment comparisons were carried out separately in each sex by DESeq2 (29) using shrinkage estimator apeglm (30) in R v4.2.1. Significance for differentially expressed genes was assumed at Benjamini-Hochberg adjusted p-value (padj) < 0.1. Normalized read counts by DESeq2 in controls and treatments were tested for gene set enrichment using Gene Set Enrichment Analysis (GSEA) v4.2.3 [build10] and C5:GO (Gene Ontology) sub-collection v7.5.1 (31). Mouse gene symbols were collapsed with human orthologs of the Molecular Signatures Database (MSigDB) v.2022.Hs.chip. Enrichment was assessed using 1000 permutations in the gene sets. Significance for enrichment was assumed at a family-wise error rate (FWER) < 0.01. Redundant GO terms from GSEA were summarized by REVIGO (http://revigo.irb.hr) (32). Gene ontology (GO) analysis for differentially expressed genes was performed using ShinyGO 0.76.3 (33). Enrichment was based on an adjusted p-value<0.05 and when no GO terms were enriched with genes with an adjusted p-value<0.05, GO terms determined from genes with a p-value<0.05 were reported, as described in (34). All RNA-Seq data are available at the Sequence Read Archive (SRA) at NCBI.

### Histology

Mice were anesthetized with Tribromoethanol (avertin) and transcardially perfused using 1x phosphate buffered saline (PBS) followed by 4% paraformaldehyde (PFA) as was done before (19). Brains were collected in PFA and incubated overnight at 4°C. Brains were transferred to 30% sucrose for 48 hours at 4°C, then frozen in OCT and cut into 30uM free-floating sections using a Leica 3050S cryostat (25), and stored at −80°C. Free-floating brain sections were processed for immunohistochemistry as before (19). Hypothalamic AgRP and POMC projections were immunostained using an AgRP (rabbit, Phoenix Pharmaceuticals INC, 1:1000) or alpha MSH (rabbit, Phoenix Pharmaceuticals INC, 1:1000) primary antibody, respectively and incubated overnight at RT. For Iba1 and GFAP, sections were incubated with Iba1 (goat, Abcam, 1:1000) or GFAP (rabbit, Millipore, 1:1000) primary antibodies overnight at RT. For pSTAT3 immunostaining, hypothalamic sections were processed by antigen retrieval (0.3% NaOH+ 1% H_2_O_2_ in PBS for 20 min; 0.3% glycine for 10 min; 0.03% SDS for 10 min; followed by normal blocking), then incubated with rabbit anti-pSTAT3 (Cell Signaling Technology) primary antibody for 48 hours at 4°C. Following several washes with PBS, sections were incubated with Alexa Fluor 488, 568 (Invitrogen 1:200) secondary antibodies for 2 hours at RT (19). Quantification of pSTAT3 positive cells in the ARC and DMH was performed at 20x magnification using Fiji-Image J as described previously (19).

### Morphometric analysis

Fluorescent images were taken with a confocal laser-scanning microscope (Zeiss LSM 800) (19) at 63X magnification. Stacked images of the ARC were analyzed using Fiji-Image J as described previously (19). For cellular morphology, skeleton and fractal analyses were performed using an established protocol (35).

### Glucose Tolerance Test

At 6 months old, following a 6-hour fast, mice received an i.p. injection of glucose (2mg/kg body weight) and blood glucose levels were measured at 0, 15, 30, 60, and 120-minutes post-injection (36).

### RNA extraction and qPCR

Total RNA was extracted from the whole hypothalamus with Trizol reagent (Gibco BRL) and 1000ng RNA was used for cDNA synthesis with the Iscript kit (Bio-Rad) as described previously (19).

### Statistical analysis

For statistical significance, data were analyzed using the Statistica software (version 14.0.015) For the GTT, a repeated measures analysis of variances (ANOVA) was used, and a two-way ANOVA analysis was conducted for the leptin signaling data set. For all ANOVA analyses, a Newman-Keuls post hoc analysis followed. For any data sets where only 2 groups of the predictor value were being compared, a two-tail unpaired Student’s *t*-test. For all statistical analyses, a 5% level of significance was set.

## Results

### Prenatal benzene exposure alters hypothalamic transcriptome in the male offspring

To evaluate the impact of prenatal benzene exposure on hypothalamic transcriptome we performed RNA sequencing on the microdissected hypothalamus from the offspring of exposed and control dams isolated at postnatal day 21 (P21) (Figure 1A). Differential gene expression analysis revealed 66 significantly changed genes when comparing gestationally benzene-exposed male offspring to controls (Fig. 1B). When comparing hypothalamic transcriptomes between benzene-exposed and control female offspring, almost no transcriptional changes were found (Fig. 1B). Among the top upregulated genes in benzene-exposed male offspring we identified genes involved in metabolic processes and directly implicated in the development of the metabolic syndrome such as *Mttp, Stk3, Ucp2, Cartpt*, and *Fgf2* (37–40) (Fig. 1B). Additionally, we identified upregulation of genes related to inflammatory response and inflammation-related pathways, such as *Tmem176a, Stxbp4, Tmem185b, Ak4, Ikzf4, and Cdk4* (41–44). Within the top downregulated genes, we identified genes associated with a neurodevelopmental function such as *Mab21l1, Ptgis, Hapln1, Ebf2*, and *Trpc3* (45–49). Downregulated genes were associated with ciliopathies in benzene-exposed male offspring, with several differentially expressed genes involved with cilium motility, assembly, and extracellular transport in benzene-exposed males (FDR<0.01) (Fig. 1C). Moreover, a gene ontology (GO) enrichment analysis of the differentially expressed genes in male offspring revealed KEGG pathway enrichment for multiple pathways, including those related to metabolic signaling pathways such as MAPK, PI3K-AKT, cAMP, and lipid metabolism (Fig. 1D). The biological and cellular compartments processes enrichment indicated enrichment for CNS developmental processes in male offspring of benzene-exposed dams (Fig. 1E, F).

**Figure 1.**
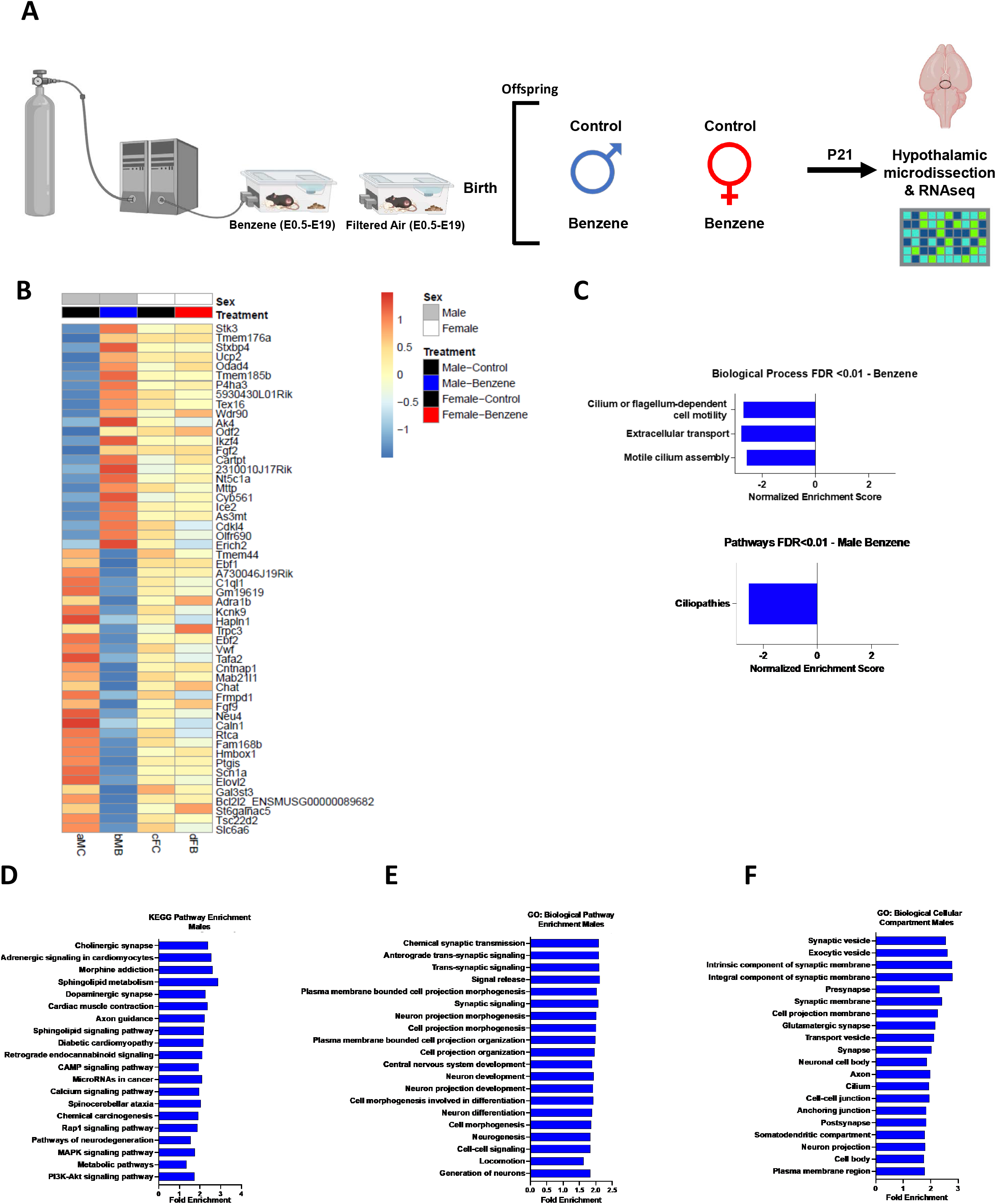
Prenatal benzene exposure alters hypothalamic transcriptome in the male offspring. (A) Experimental design of maternal benzene exposure and RNAseq of the hypothalamus of male and female offspring (n=3-5 per litter, from 3-4 litters/condition) (B) Average differentially expressed genes (DEGs) of both male and female offspring P21 hypothalamus. (C) Biological process and pathways of the top DEGs in male offspring (FDR< 0.01). (D) KEGG pathways enrichment of DEGs in male offspring (p<0.05) (E) Biological pathway enrichment of male offspring DEGs (p<0.05) (F) Enrichment of cellular compartments in DEGs of male offspring (p<0.05).

### Prenatal benzene exposure alters hypothalamic orexigenic and anorexigenic projections in male offspring

We next assessed the impact of prenatal benzene exposure on the hypothalamic orexigenic and anorexigenic pathways, known for their role in the control of satiety signals and metabolic regulation (21). We analyzed the immunoreactivity of agouti-related protein (AgRP) and α-melanocyte-stimulating hormone (α-MSH) (pro-opiomelanocortin precursor) containing fibers in the paraventricular nucleus of the hypothalamus (PVH) at P21 (Fig. 2A, B). Quantification of the fiber density in the anterior PVH revealed significant decreases in both AgRP and α-MSH fiber densities in benzene-exposed male offspring compared to controls (Fig. 2C, D). There were no significant differences in AgRP and α-MSH fiber densities between prenatally benzene-exposed or control female offspring.

**Figure 2.**
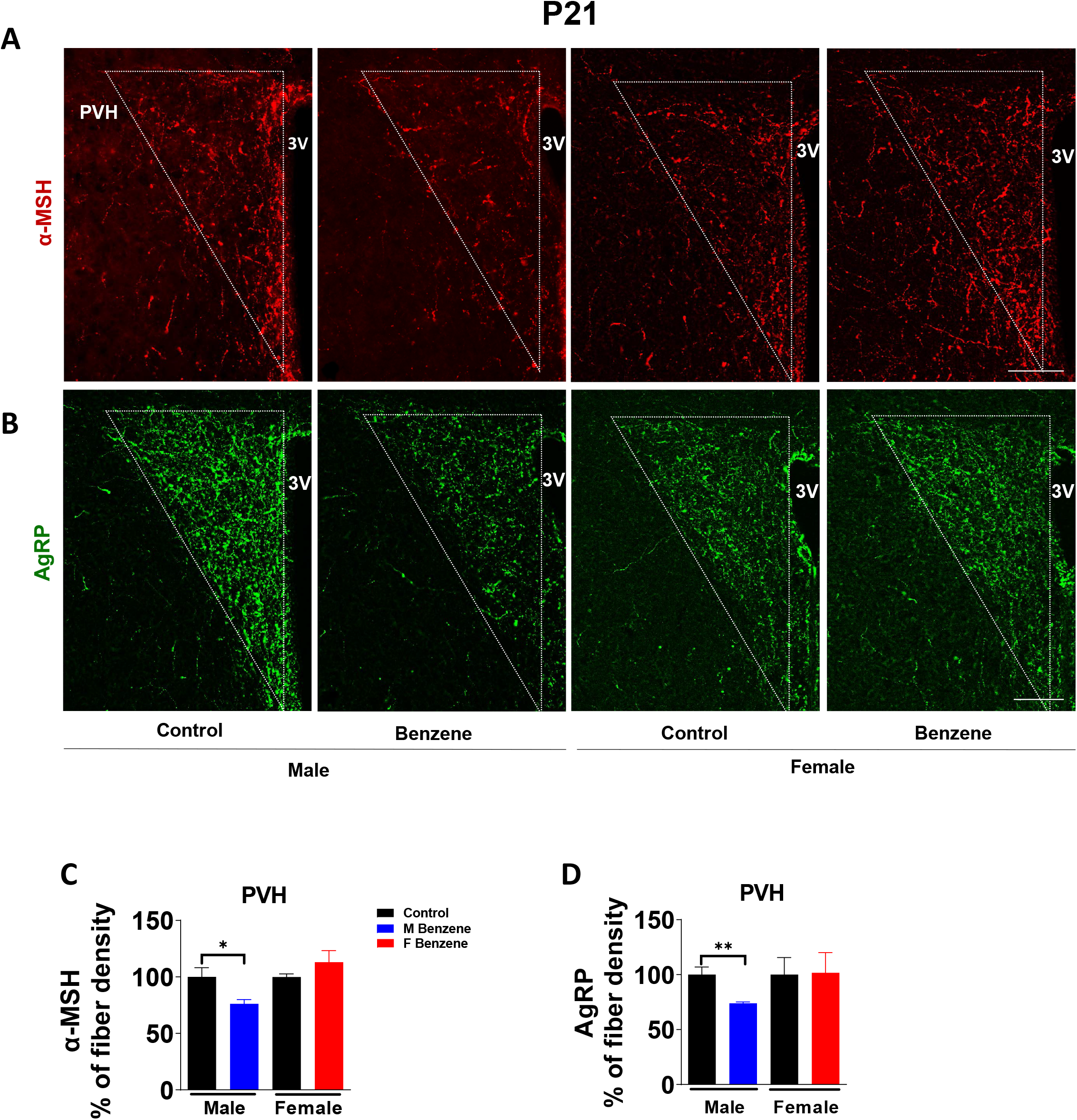
Prenatal benzene exposure alters hypothalamic orexigenic and anorexigenic projections in male offspring. (A) Representative images of neuronal projections in the PVH identified by immunofluorescent detection of alpha-melanocyte-stimulating hormone (α-MSH) in P21 offspring. (B) Representative images of neuronal projections in the PVH identified by immunofluorescent detection of agouti-related protein (AgRP) in P21 offspring. (C) Quantitation of α-MSH fiber density in male and female offspring at 20x magnification (n= 3-5 per group). (D) Quantitation of AgRP fiber density in male and female offspring at 20x magnification (n= 3-5 per group). Data were expressed as the mean ± SEM and analyzed by t-test (*= vs control; *p<0.05, **p<0.01).

### Prenatal benzene exposure alters hypothalamic leptin signaling in male offspring

The proper development of a hypothalamic nutrient circuitry is essential for leptin-responsive neurons and leptin signaling (50). Leptin binding to LepRb activates an associated Jak2 tyrosine kinase, thereby promoting the phosphorylation of LepRb and the recruitment and tyrosine phosphorylation of Stat3 (signal transducer and activator of transcription-3) (51). To evaluate hypothalamic leptin responses, we measured leptin-stimulated accumulation of pStat3 in the offspring hypothalamus, which is known to be impaired in states associated with diminished leptin action (52). Acute leptin treatment led to a similar increase in pStat3 levels in the arcuate nucleus (ARC) of the hypothalamus of both male and female offspring (Fig. 3A, B, D). However, a significant decrease (p < 0.001) in pStat3 positive cells was observed in the dorsomedial hypothalamus (DMH) only in benzene-exposed male offspring, but not females (Fig. 3C). Thus, prenatal benzene exposure alters hypothalamic leptin signaling of male offspring, affecting the organization of neural circuits that regulate energy homeostasis and glucose metabolism, which may contribute to sex-specific metabolic alterations previously observed in this model (8).

**Figure 3.**
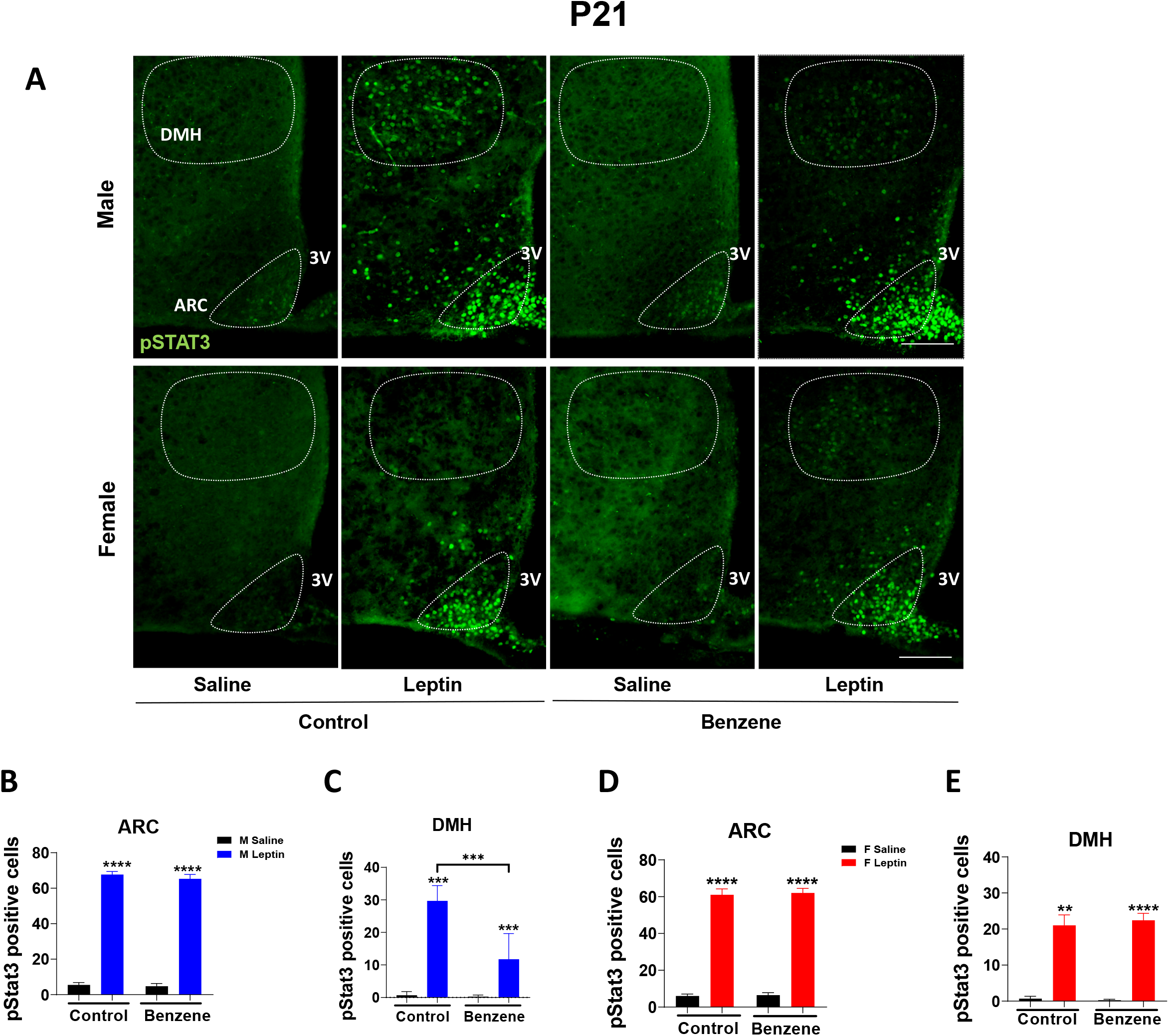
Prenatal benzene exposure alters hypothalamic leptin signaling in male offspring. (A) Representative images of phosphorylated signal transducer and activator of transcription −3 (pSTAT3) positive cells in the arcuate nucleus (ARC) and dorsomedial hypothalamic nucleus (DMH) of P21 offspring. Quantification of pSTAT3+ cells in the ARC identified by immunofluorescence in male (B) and female (D) offspring at 20x magnification (n=3-5 per group). Quantification of pSTAT3+ cells in the DMH of male (C) and female (E) P21 offspring. (F) Relative mRNA expression of epigenetic modifying genes from offspring P5 hypothalamus of both sexes. Data were expressed as the mean ± SEM and analyzed by *t*-test if necessary to compare between only two groups of predictor variable or Two-way ANOVA followed by Newman-Keuls post hoc analysis, (* = *vs* control; *p < 0.05; **p < 0.01; ***p < 0.001; ****p < 0.0001).

### Prenatal benzene exposure induces hypothalamic microgliosis in the offspring

To assess the effect of prenatal benzene exposure on hypothalamic inflammation, we analyzed the numbers and cellular morphology of microglia and astrocytes in the offspring at P21. We found a significant increase in the microglia-specific ionized calcium-binding adaptor molecule 1 (Iba1) marker, in the ARC of benzene-exposed male but not female offspring (Fig. 4A, E). Interestingly, the analysis of microglial morphology revealed increased branch length and endpoints in both male and female benzene-exposed offspring, along with increased density in male mice (Fig. 4C, F, I, J). Importantly, there were no significant alterations in astrocyte numbers or cellular morphology between control and benzene-exposed offspring of both sexes (Fig. 4B, D, G, H, K, L). We further found an overall reduction in glial gap junction protein CX30 immunoreactivity in benzene-exposed male offspring (Supplementary Fig. 1A and B). Since CX30 is highly associated with astrocyte structure and function (53), we analyzed the colocalization of CX30 with both microglia and astrocytes (Supplementary Fig. 1A-D). However, as measured by the Pearson correlation coefficient, we observed no differences in the colocalization of CX30 with astrocytes or microglia in benzene-exposed offspring of either sex (Supplementary Fig. 1C, D).

**Figure 4.**
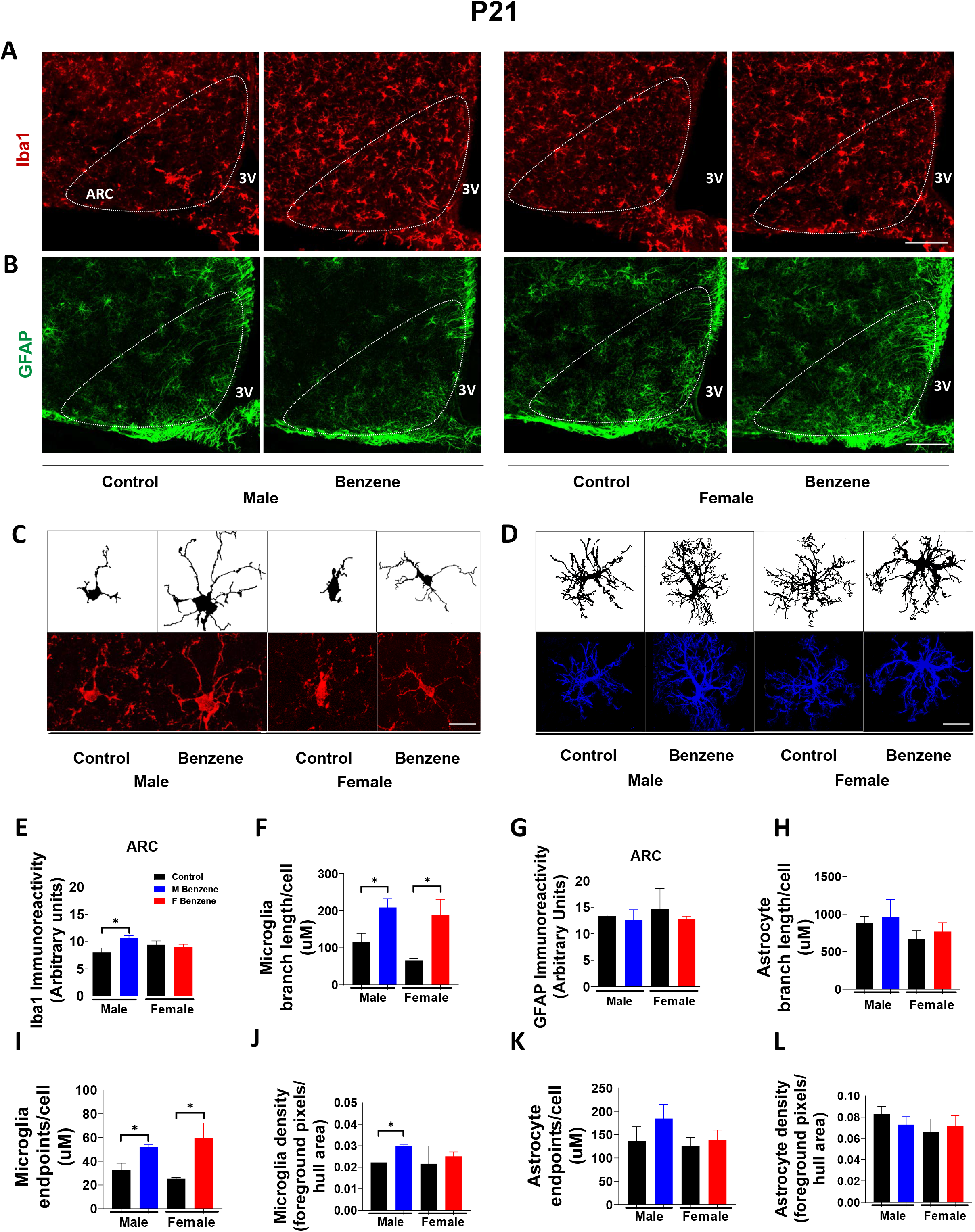
Prenatal benzene exposure induces hypothalamic microgliosis in the offspring. (A) Representative images of microglia identified by immunofluorescent detection of Ionized calcium-binding adaptor molecule 1 (Iba1) in P21 ARC at 10x magnification. (B) Representative images of astrocytes identified by immunofluorescent detection of glial fibrillary acidic protein (GFAP) in P21 ARC at 10x magnification. (C) Microglial morphology in P21 offspring of both sexes. (D) Astrocyte morphology in P21 offspring of both sexes. (E) Quantitation of Iba1 immunoreactivity (n=5 per group). (G) Quantitation of GFAP immunoreactivity (n=3-5 per group). (F; H-L) Measurement of microglial and astrocyte branch length, endpoints, and density in P21 offspring by skeleton analysis (see Methods). Data were expressed as the mean ± SEM and analyzed by *t*-test (*= vs control; *p<0.05, **p<0.01, ***p<0.001).

### Benzene exposure during lactation does not exacerbate hypothalamic gliosis in the offspring

We next explored if subsequent benzene exposure in early postnatal life, during lactation, when the hypothalamus is still developing in rodents (22), would exacerbate the observed hypothalamic inflammation in the offspring. Surprisingly, additional exposure to benzene during lactation did not further increase microglia numbers in the male offspring compared to controls (Fig. 5B, D), emphasizing the importance of the developmental stages on microglia sensitivity and activation. Importantly, exposure to benzene during lactation did not affect astrogliosis in the offspring of both sexes (Fig. 5C, E).

**Figure 5.**
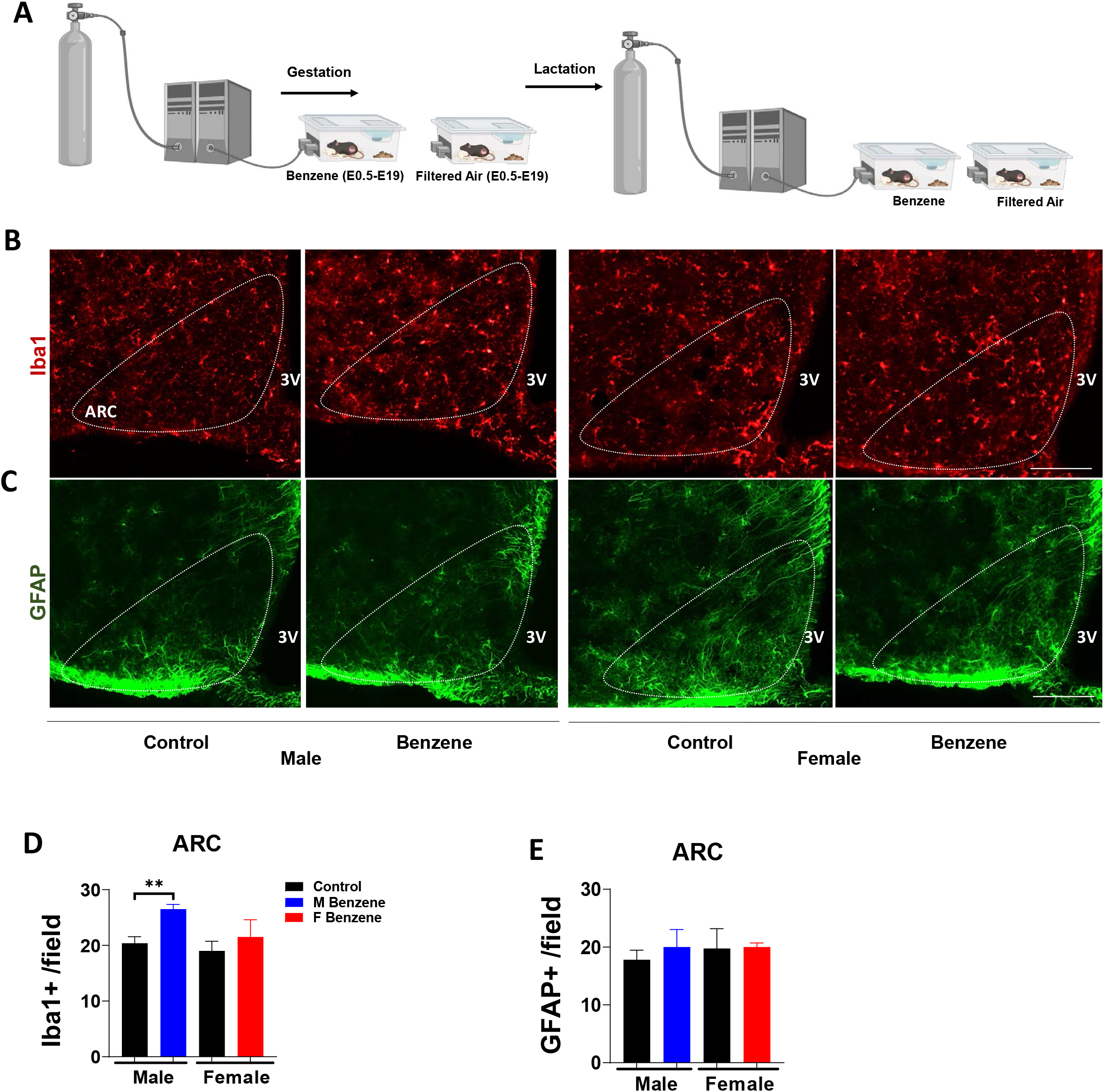
Benzene exposure during lactation does not exacerbate hypothalamic gliosis in the offspring. (A) Representative images of microglia identified by Iba1 immunofluorescence in male and female P21 offspring ARC. (B) Representative images of astrocytes identified by GFAP immunofluorescence in male and female P21 offspring ARC. Quantification of microglia (C) and astrocytes (D) in P21 offspring of both sexes at 10x magnification (n=4-5 per group). Data were expressed as the mean ± SEM and analyzed by *t*-test (*= vs control; ***p<0.001).

### Prenatal benzene exposure predisposes HFD-fed offspring to susceptibility to metabolic alterations in adulthood

Five months of high-fat diet (HFD) challenge from 4 months of age till 9 months of age did not magnify weight gain or fat/lean mass among male or female offspring of exposed mothers (Fig. 6A-C and Supplementary Figure 2). However, the HFD challenge significantly impaired glucose tolerance in both male and female offspring of benzene-exposed dams (Fig. 6D and E), suggesting that maternal benzene exposure contributes to diet-induced hyperglycemia triggered by HFD. Interestingly, this effect was more profound in female offspring (ANOVA p<0.05 control vs benzene) (Fig. 6E). HFD-fed male offspring of benzene-exposed dams demonstrated a significantly higher number of immunostained Iba1^+^ microglia (p<0.01) and GFAP^+^ astrocytes (p<0.05) in the hypothalamus compared to HFD controls (Fig. 6F and G). Importantly, female offspring of benzene-exposed dams demonstrated increased numbers of GFAP^+^ astrocytes similar to male benzene offspring (Fig. 6I), suggesting that maternal benzene exposure eliminated the protective gender effect associated with diet-induced hypothalamic inflammation (54). Together, our data demonstrate long-term hypothalamic changes as a result of maternal exposure to benzene during pregnancy, changes which worsen specifically the offspring’s ability to cope with the HFD challenge.

**Figure 6.**
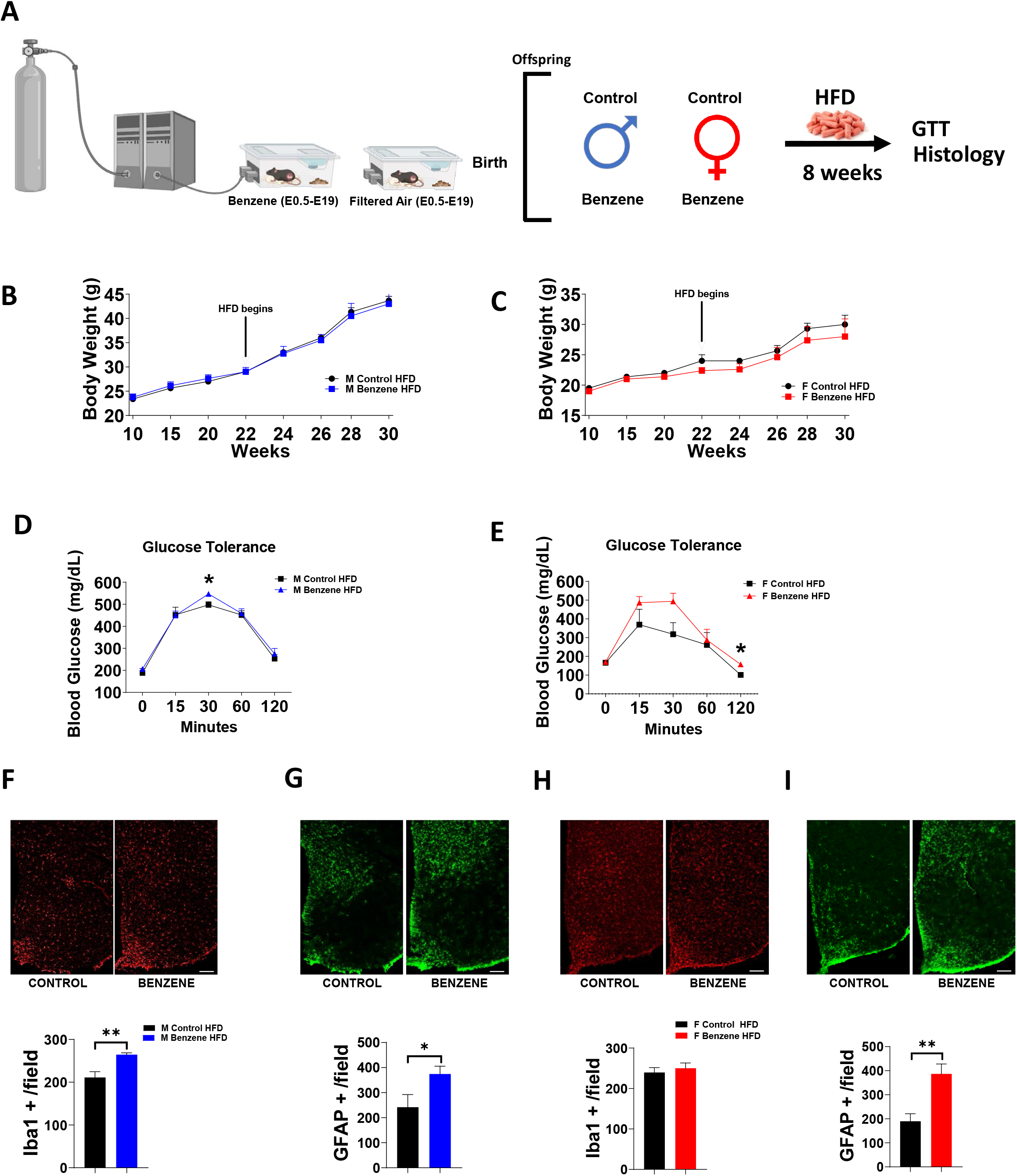
Prenatal benzene exposure predisposes HFD-fed offspring to susceptibility to metabolic alterations in adulthood. (A) Timeline of experimental design. (B) Body weight (g) in adult HFD males (B) and females (C). Glucose tolerance test in 6-month-old HFD males (D) and females (E). Quantification of Iba1+ cells in the MBH of male (F) and female (H) 6-month-old offspring. Quantification of GFAP+ cells in the MBH of male (G) and female (I) 6-month-old offspring. Data were expressed as mean ± SEM (n=3-4 per group) and analyzed by *t*-test if necessary to compare between only two groups of predictor variable or repeated measures ANOVA followed by Newman-Keuls post hoc analysis, (*= vs control; *p<0.05).

## Discussion

There is mounting epidemiological evidence for an increased risk of metabolic disease in the offspring of mothers who smoke (12, 55). Notably, cigarette smoke is the primary source of non-occupational benzene exposure in the US population (56, 57). We have recently demonstrated that *in-utero* exposure of pregnant female mice to benzene at concentrations relevant to smoking leads to altered energy homeostasis and glucose intolerance in the offspring, with a stronger effect in the male offspring (8). Here we demonstrate for the first time that a window of vulnerability exists during gestational development where the hypothalamus is especially sensitive to such exposure, causing irreversible damage to the hypothalamic nutrient and metabolic circuits, neuroinflammation, and alterations in the hypothalamic transcriptome, especially in male offspring. Notably, prenatal but not the post-natal exposure window was critical in hypothalamic sensitivity to the benzene exposure, indicating the importance of prenatal environmental exposure for hypothalamic development. Regardless, both male and female offspring born to benzene-exposed dams exhibit exacerbated hyperglycemia and hypothalamic inflammation when challenged with HFD, providing a causal relationship between *in-utero* benzene exposure and metabolic disease susceptibility.

Only benzene-exposed male but not female offspring exhibited changes in the hypothalamic transcriptional profile, mostly affecting the biological functions associated with brain developmental processes, cilia function, and extracellular transport. Early observations show the correlation between metabolic disorders and neuronal primary cilia function (58, 59), while ablation of ciliary genes results in hyperphagia-induced obesity and hyperglycemia (60). Indeed, we detected the upregulated genes related to metabolic processes including PI3K-Akt signaling cascade and lipid homeostasis. Some of these genes, for example, *Stxbp4, Stk3*, and *Mttp* were found to be involved in pathways related to diabetes risk, diabetes complications, or risk of cardiovascular disease in population studies (61–63). Interestingly, changes in Cart and Fgf2 levels has been associated with hypothalamic-pituitary-adrenal axis activity and leptin signaling (64, 65), linking these alterations with hypothalamic postnatal hormonal regulation in benzene-exposed male offspring. Another upregulated gene, *Ucp2*, was shown to induce hypothalamic microglia inflammation in obesity (66), suggesting dysregulation of hypothalamic microglia in the male benzene-exposed offspring. Importantly, metabolic changes co-occur with changes in genes that affect CNS development and function, highlighting the susceptibility of the developing hypothalamus to environmentally-induced stress. These early changes in the genes profile in the hypothalamus might explain the stronger predisposition to the peripheral metabolic imbalance in benzene-exposed male offspring compared to females (8).

Prenatal benzene exposure led to a rapid and long-lasting decrease of the opposing catabolic α-MSH and anabolic AgRP projections to the PVN exclusively in the male offspring. The diminished ARC-PVN pathway may reflect, or contribute to, the early decrease in hypothalamic leptin sensitivity and developmental decline of gap junction CX30 reported in this study as well as in previously published models of gestational diet-induced obesity (DIO) (27, 67–70). Similarly, alterations in the organization of hypothalamic neural circuits controlling appetite may contribute to the metabolic effects observed in adult male offspring. In support, adult male offspring from benzene-exposed dams show severe hyperglycemia and hyperinsulinemia with elevated β-cells mass (8).

Disruption of hypothalamic development by neuroinflammation plays a key role in the pathogenesis of the metabolic disease, as inflammation is one mechanism that can induce longterm neurodevelopmental changes in the CNS (71). In support, prenatal benzene exposure induced upregulation of hypothalamic genes related to the inflammatory and immune response in male offspring. For example, *Ak4* was shown to promote inflammatory gene expression and secretion of inflammatory cytokines from activated macrophages (42), *Ikzf4* regulates immune cell development and cytokines signaling (72), while *Cdk4* is a major target for hypothalamic NF-kβ inflammatory pathway and a target for central anti-obesity treatment (73). Upregulation of inflammatory genes might contribute to hypothalamic microgliosis observed in male offspring. Interestingly, no significant changes were observed in astrocyte numbers or morphology in the offspring of benzene-exposed dams. Moreover, the combined prenatal and postnatal benzene exposure during the lactation had no significant effect on astrogliosis and did not promote further microglial alterations in male offspring. In models of maternal DIO, the proportion of proliferating astrocytes was significantly higher in the ARC of the DIO-offspring compared to control offspring (74). These findings highlight the differences between nutritional and environmental stressors on adverse neurodevelopmental outcomes. Collectively, maternal *in-utero* benzene exposure caused enduring changes in the microglia of adult offspring which might affect the offspring’s susceptibility to metabolic alterations.

The weak impact of prenatal/postnatal benzene exposure in female offspring in our study is consistent with previous reports showing that male offspring are more likely to develop behavioral, cognitive, and epigenetic brain changes following exposure to a perinatal HFD or diesel exhaust (71, 75, 76). For example, prenatal air pollution exposure predisposed male but not female offspring to increased neuroinflammation and susceptibility to DIO (71). Maternal smoking during pregnancy led to lower birth weight and small-for-gestational-age male infants (77). Moreover, prenatal exposure to diesel exhaust PM_2.5_ programs the homeostatic regulation of glucose metabolism in adult male offspring (78). Our data support our previous work showing a male predisposition to metabolic imbalance, since only male offspring from benzene-exposed dams were glucose intolerant and insulin resistant at 4 months of age (8). By 6 months of age, both male and female offspring displayed glucose and insulin intolerance, associated with elevated expression of hepatic gluconeogenesis and inflammatory genes (8), suggesting that it takes longer for females to develop aberrations in metabolic phenotype. Interestingly, only adult males exposed to benzene exhibit severe metabolic imbalance associated with hypothalamic inflammation and endoplasmic reticulum (ER) stress, suggesting sex-specific protection from the adverse consequences of benzene exposure. Sex hormones have been linked to sex differences in metabolic responses as well as in microglia and astrocytes transcriptome heterogeneous (79–81). Further investigation will be required to determine how sex hormones contribute to differential vulnerability between males and females to perinatal/post-natal benzene exposure across the lifespan of male and female offspring.

Despite similar weight gain on HFD, both male and female offspring of benzene-exposed dams showed exacerbated hyperglycemia and hypothalamic gliosis following HFD feeding at 6 months of age. The effect of HFD on female offspring of benzene-exposed dams is especially interesting since females failed to show major hypothalamic deficits at a young age. Although, increased microglial activation following HFD challenge in adulthood, suggests that female microglia were primed during prenatal benzene exposure, as indicated by changes in hypothalamic microglia morphology in these mice. Future studies are needed to assess the impact of microglia changes in response to the HFD challenge to define the mechanisms underlying benzene-induced prenatal metabolic programming.

Overall, our study provides causal evidence that maternal benzene exposure impairs the metabolic health of the child through hypothalamic changes during pregnancy. Maternal environmental benzene exposure can occur as a result of smoking, exposure to secondhand smoke, as well as due to living in a high-pollutant urban environment. As rates of environmental exposures continue to rise, the current findings may have implications for the future incidence of metabolic disease in offspring beyond middle age.

## Abbreviations

T2DM: Type 2 Diabetes Mellitus
VOCs: volatile organic compounds
ARC: Arcuate nucleus
DMH: Dorsomedial hypothalamus
AgRP: agouti-related protein
α-MSH: α-melanocyte-stimulating hormone
POMC: pro-opiomelanocortin
pSTAT3: phosphorylated signal transducer and activator of transcription 3
Iba1: ionized calcium binding adaptor protein 1
GFAP: glial fibrillary acidic protein
HFD: high-fat diet

## Author Contributions

LS, SS, LKD, LS, MS, and HJ carried out the research and reviewed the manuscript. PF, JEPM, and EG assisted in data analysis. AL analyzed the data and reviewed the manuscript. MS designed the study, analyzed the data, wrote the manuscript, and is responsible for the integrity of this work. All authors approved the final version of the manuscript.

## Acknowledgments

This project was supported by American Diabetes Association grant #1-lB-IDF-063, CURES Center Grant (P30 ES020957), NIEHS R01ES033171 and CLEAR P42ES030991 for MS. LK was further supported by NIH (5T32GM142519-02). The author(s) thank the Van Andel Genomics Core for providing RNA-sequencing facilities and services.

## Disclosure

No conflict of interest, financial or otherwise, is declared by the authors.

## Data Availability Statement

The data that support the findings of this study are available from the corresponding author upon reasonable request.

## Notes

### Competing Interest Statement

The authors have declared no competing interest.

