## Supplementary Material for "Prenatal benzene exposure alters offspring hypothalamic development predisposing to metabolic disease in later life"

### Supplementary Figure 1

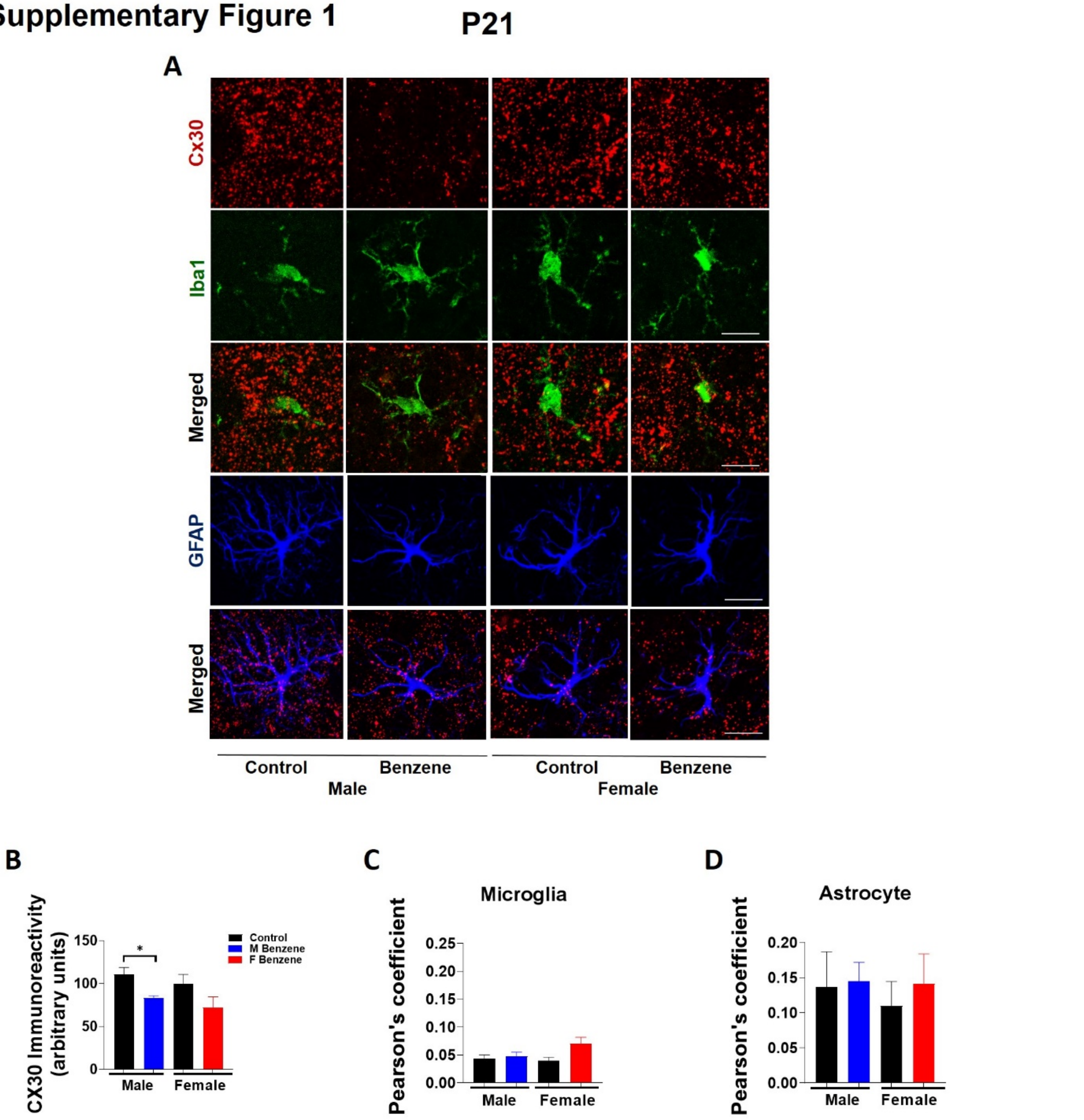


**Supplementary Figure 1. Prenatal benzene exposure reduces gap junction in male offspring.** Representative images of colocalization between gap junctions and microglia (A) identified by immunofluorescent detection of connexin 30 (CX30) and Iba1, or astrocytes (C) identified by GFAP at 63X magnification in P21 ARC. (B) Quantification of total CX30 immunoreactivity in the ARC (n=3-4 per group). (C) Quantitation of Pearson's colocalization coefficient between CX30 and microglia in the ARC. (D) Quantitation of Pearson's colocalization coefficient between CX30 and astrocytes in the ARC. Data were expressed as the mean ± SEM and analyzed by *t*-test (*= vs control; *p<0.05).

### Supplementary Figure 2

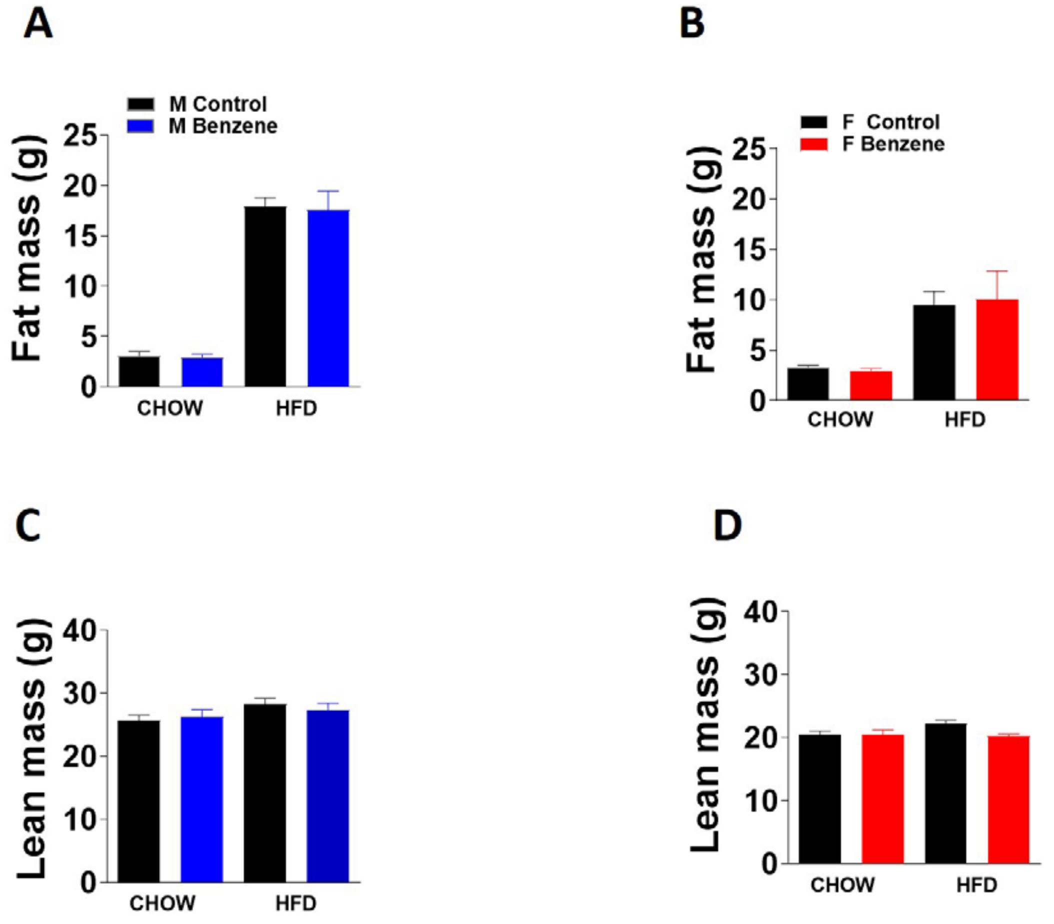


**Supplementary Figure 2. Prenatal benzene exposure does not increase fat or lean mass in HFD-fed offspring.** Fat mass of 9-month-old male (A) and female (B) offspring. Lean mass of 9-month-old male (C) and female (D) offspring. (n=3-5 per group). Data were expressed as the mean ± SEM and analyzed by *t*-test (*= vs control; *p<0.05).
